# Drug discovery for heart failure targeting myosin-binding protein C

**DOI:** 10.1101/2023.04.03.535496

**Authors:** Thomas A. Bunch, Piyali Guhathakurta, Andrew R. Thompson, Victoria C. Lepak, Anna L Carter, Jennifer J. Thomas, David D. Thomas, Brett A. Colson

**Affiliations:** Department of Cellular & Molecular Medicine, University of Arizona, Tucson Arizona 85724; Department of Biochemistry, Molecular Biology, and Biophysics, University of Minnesota, Minneapolis, MN 55455; Photonic Pharma LLC, Minneapolis, MN, USA

**Author notes:** Equal contribution (Thomas A. Bunch, Piyali Guhathakurta). Co-senior authors (David D. Thomas, Brett A. Colson).

**Keywords:** actin, cardiac muscle, cardiac myosin-binding protein C (cMyBP-C), contractile proteins, phosphorylation, protein kinase A (PKA), fluorescence lifetime (FLT), high throughput screen (HTS), fluorescence resonance energy transfer (FRET), site-directed spectroscopy

## Abstract

Cardiac MyBP-C (cMyBP-C) interacts with actin-myosin to fine-tune cardiac muscle contractility. Phosphorylation of cMyBP-C, which reduces binding of cMyBP-C to actin or myosin, is often decreased in heart failure (HF) patients, and is cardioprotective in model systems for HF. Therefore, cMyBP-C is a potential target for HF drugs that mimic phosphorylation and/or perturb its interactions with actin or myosin. We labeled actin with fluorescein-5-maleimide (FMAL), and the C0-C2 fragment of cMyBP-C (cC0-C2) with tetramethyl rhodamine (TMR). We performed two complementary high-throughput screens (HTS) on an FDA-approved drug library, to discover small molecules that specifically bind to cMyBP-C and affect its interactions with actin or myosin, using fluorescence lifetime (FLT) detection. We first excited FMAL and detected its FLT, to measure changes in fluorescence resonance energy transfer (FRET) from FMAL (donor) to TMR (acceptor), indicating binding and/or structural changes in the protein complex. Using the same samples, we then excited TMR directly, using a longer wavelength laser, to detect the effects of compounds on the environmentally sensitive FLT of TMR, to identify compounds that bind directly to cC0-C2. Secondary assays, performed on selected modulators with the most promising effects in the primary HTS assays, characterized specificity of these compounds for phosphorylated versus unphosphorylated cC0-C2 and for cC0-C2 versus C1-C2 of fast skeletal muscle (fskC1-C2). A subset of identified compounds modulated ATPase activity in cardiac and/or skeletal myofibrils. These assays establish feasibility for discovery of small-molecule modulators of the cMyBP-C-actin/myosin interaction, with the ultimate goal of developing therapies for HF.

## Introduction

Cardiomyopathies are the most common and severe inherited disorders that are associated with significant adverse outcomes such as heart failure (HF), arrhythmias, and sudden cardiac death (1). Hypertrophic cardiomyopathy (HCM) results from mutations in 11 or more different sarcomeric genes (2) and is characterized by hypertrophy of the left ventricle and the interventricular septum, typically reducing ventricular chamber volume and causing myocyte and myofibrillar disarray. In HCM, the heart typically becomes enlarged, hypercontractile, and unable to relax effectively, but the clinical manifestations of the disease are quite variable (3). The wide spectrum of functional perturbations induced by the different HCM mutations suggests that many different pathways lead to the HCM phenotype, so it is difficult to establish a prognosis based on the mutation (4). Dilated cardiomyopathy (DCM) is the leading cause of heart transplantation and is mainly characterized by cardiac hypocontractility and enlargement of the ventricular chambers, leading to progressive HF (5,6).

The primary cause of HCM is most often a mutation in one of several sarcomeric proteins, including cMyBP-C, myosin, actin, troponin, tropomyosin, leiomodin, and titin. cMyBP-C is a sarcomeric regulatory protein that modulates muscle contraction/relaxation by interacting with both actin and myosin (Figure 1). Numerous studies have demonstrated that increasing or decreasing protein kinase A (PKA)-mediated phosphorylation of cMyBP-C allows for tuning cardiac contraction and relaxation through phosphorylation-sensitive interactions with actin and myosin. Recent work by us (7) and others (8) suggests that phosphorylation induces a structural rearrangement in the N-terminal domains of cMyBP-C, which reduces its binding to actin and myosin, affecting muscle contraction and relaxation. Decreases in phosphorylation of cMyBP-C have been observed in HF patients, including those affected by mutations in sarcomeric proteins other than cMyBP-C (9,10). Therefore, targeting cMyBP-C with drugs that mimic phosphorylation and/or perturb its interactions with actin or myosin is a promising approach to improving cardiac muscle function in HF and cardiomyopathy.

**Figure 1.**
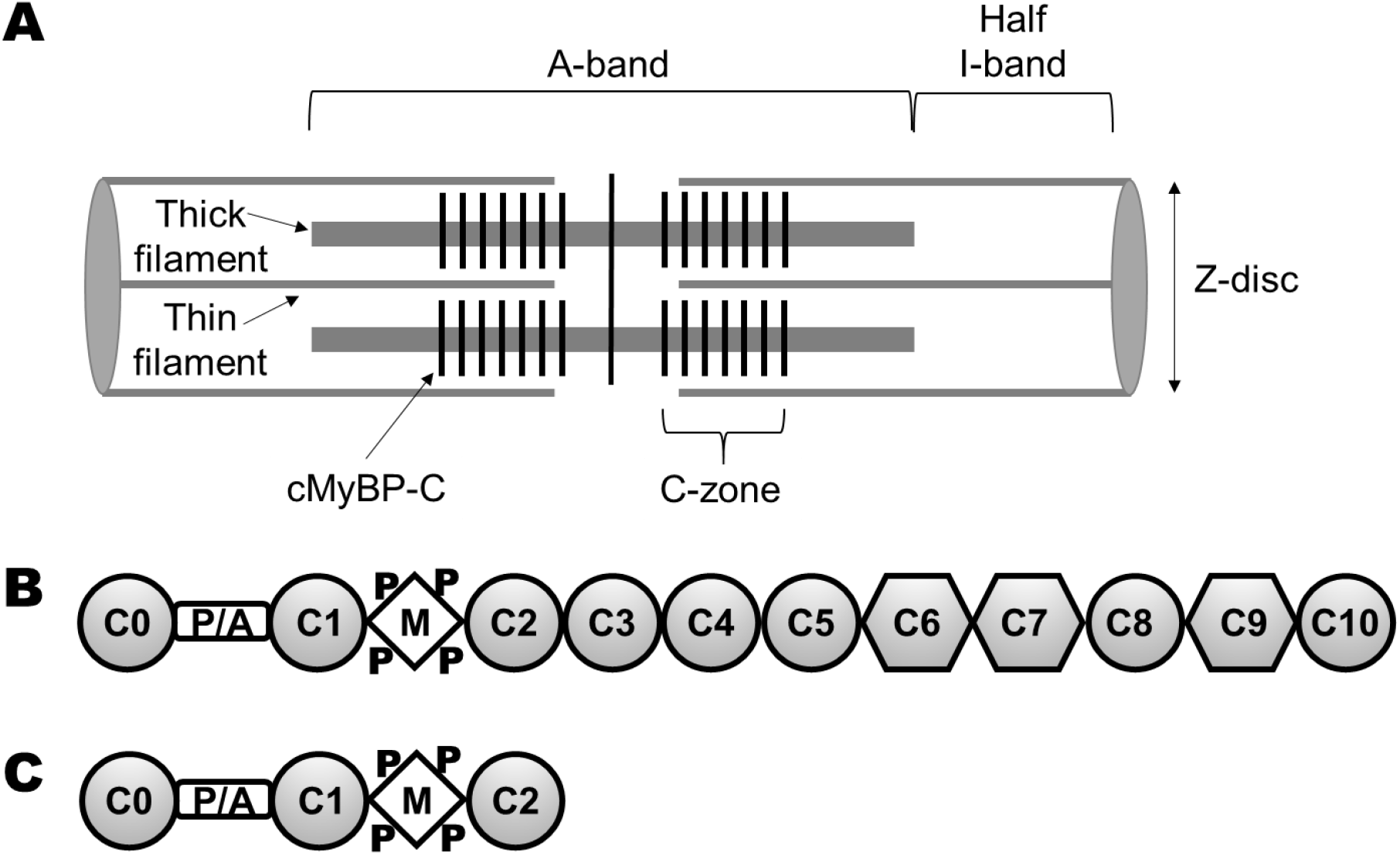
cMyBP-C organization in the sarcomere. (A) The sarcomere spans from Z-disc to Z-disc with the A-band containing thick filaments and the I-band containing actin filaments. Force is generated by myosin and actin in the thin/thick filament overlap portion of the A-band. cMyBP-C molecules (black vertical stripes) are anchored to the thick filament and present in the C-zones toward the center of the A-band. The C-zones overlap with thin filaments (except at very long sarcomere lengths). (B) Full-length cMyBP-C domains C0 through C10. Ig-like domains are shown as circles and fibronectin type-III domains are shown as hexagons. (C) N-terminal domains C0 through C2 (C0-C2), containing the proline alanine rich linker (P/A) and the M-domain (M) that contains phosphorylation sites (P) (adapted from (13)).

Small molecule modulators targeting various sarcomeric proteins have been developed in recent years, with some progressing to clinical trials. Omecamtiv mecarbil (OM), a selective cardiac myosin activator (11) (developed by Cytokinetics, Inc.), and mavacamten (Mava), a selective β- cardiac myosin inhibitor (developed by MyoKardia, Inc) (12), are so far the most promising examples. However, cardiomyopathies are diverse, and a variety of modulators targeting other sarcomeric proteins is desirable in order to provide adequate treatment options for patients.

We previously reported a HTS assay, based on the detection of fluorescence lifetime (FLT) with subnanosecond time resolution, and identified three unique cMyBP-C-binding compounds (13). However, these compounds were subsequently found to not be specific for cMyBP-C, as they inhibit fast skeletal MyBP-C (fskMyBP-C) interactions with actin (Fig. S1) and bind to both slow skeletal (ssk) and fsk MyBP-C N-terminal domains (Fig. S2). In the current study, we have extended and enhanced our previous approach, developing and performing two complementary high-throughput FLT-based screens to identify compounds that are muscle-type specific. We labeled actin with fluorescein-5-maleimide (FMAL) and the C0-C2 fragment of cMyBP-C with tetramethylrhodamine (TMR), used as an acceptor. We first performed HTS to identify compounds that influence the interaction between actin and cC0-C2, using FLT-detected measurement of fluorescence resonance energy transfer (FRET) from donor-labeled actin to acceptor-labeled cC0-C2. Donor (FMAL) on actin was excited at 473 nm, and FRET to the acceptor (TMR) on cC0-C2 was measured using a high-precision, high-speed fluorescence lifetime plate reader (FLTPR) (14). We then performed a second screen, FLT-TMR, using the same samples in the same plate, to identify compounds that bind directly to cC0-C2, whether or not they affect its actin binding. Here the TMR on cC0-C2 was excited at 532 nm using a different laser in the same FLTPR, and FLT changes due to the presence of compound identified cC0-C2-binding structural modulators. These compounds are potential effectors for cC0-C2 interactions with myosin. Using these two approaches, we screened a 2,684-compound Selleck library of FDA-approved compounds (Selleckchem) with the goal of identifying cMyBP-C-binding compounds that modulate its activities in the sarcomere.

We established additional and reliable biochemical and structural secondary assays, which were lower throughput than the primary FLT assays, but rapid and precise enough to test whether the hit compounds from the primary HTS screens are specific for cMyBP-C versus fskMyBP-C. Differences in binding to phosphorylated versus non-phosphorylated cMyBP-C were also measured. Finally, we tested the effect of these compounds on the ATPase activities of skeletal and cardiac myofibrils, to determine whether these hits modulate cardiac and skeletal function. A clear pathway to finding the lead compounds targeting cMyBP is shown in Figure 2.

**Figure 2:**
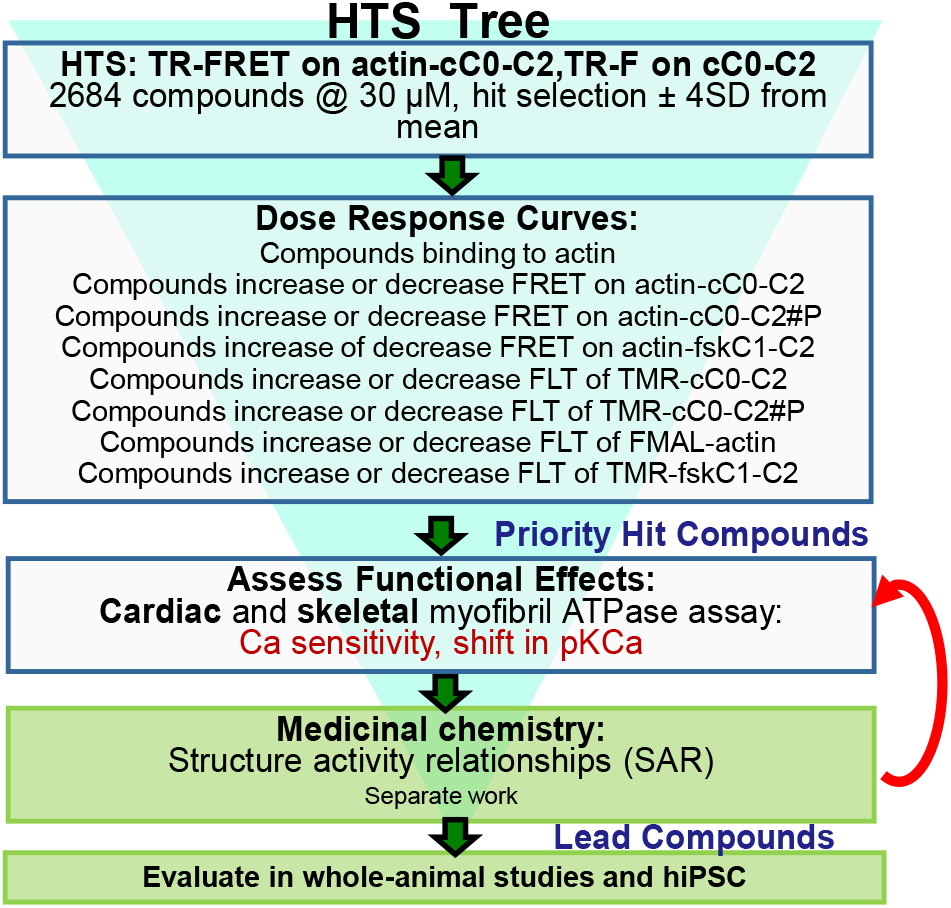
HTS process (funnel) for finding compounds targeting cMyBPC.

The success of these high-throughput primary and secondary assays, demonstrated below, gives us confidence that this approach can be expanded to much larger libraries to identify more compounds to serve as starting points for the medicinal chemistry that will probably be necessary to increase specificity and drug potency for the treatment of HF.

## Results

### FLT-FRET of Actin-cC0-C2/fskC1-C2 biosensor

Using a FLT-detected cC0-C2-actin binding assay, we previously identified the first three compounds (13) that bind to cC0-C2 and inhibit its interactions with actin (13,15). However, these compounds were subsequently found to not be specific for cMyBP-C, as they inhibit fast skeletal MyBP-C (fskMyBP-C) interactions with actin (Fig. S1). The effects of these compounds on cC0-C2 binding were confirmed with a novel FLT-detected FRET assay, in which a donor (FMAL) was attached to actin, and an acceptor (TMR) was attached to cC0-C2. Binding was monitored by decrease in FLT of FMAL (increase in FRET) (Figure 3). The FRET response (∼45% decrease in FLT) was much larger than the FLT change detected in the absence of acceptor (∼7%) (13), confirming that the primary effect of the compound was a structural change in the complex between actin and cC0C2. Preliminary trials, using the same concentration (1 μM) of labeled F-actin plus 2 μM of TMR-cC0-C2 resulted in clogging of the dispensing pins of the multidrop liquid dispenser. Therefore, we reduced the concentration of donor (FMAL-actin) and acceptor (TMR-cC0-C2) to 0.25 μM and 0.5 μM, respectively. FLT-FRET binding curves using these concentrations of FMAL-F-actin and TMR-cC0-C2 indicated that these levels are suitable to use for screening for compounds that decrease or increase binding. Binding curves for PKA-phosphorylated TMR-cC0-C2 (cC0-C2#P) and TMR-fskC1-C2 were generated, to be used in secondary assays to test the specificity of identified compounds for cMyBP-C that is or is not phosphorylated (Figure 3).

**Figure 3.**
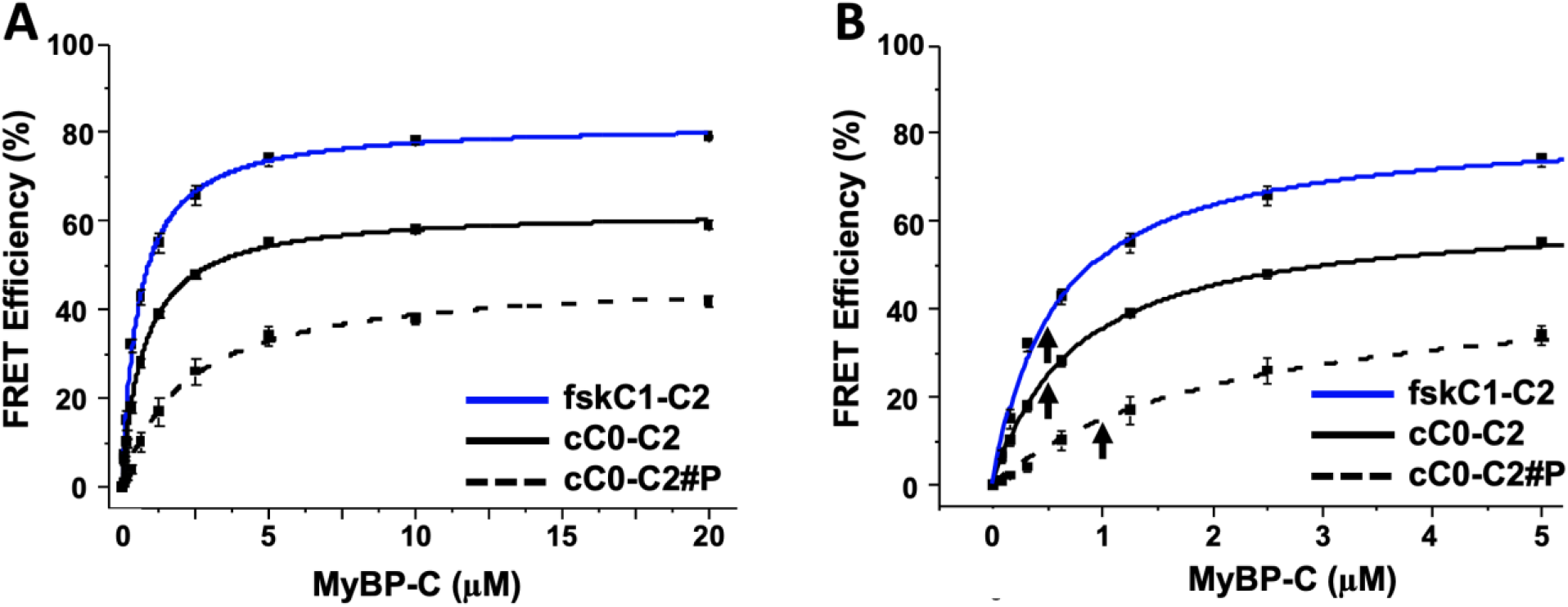
Actin-cMyBPC FLT-FRET assay. (A)FLT-FRET-based binding curve of 0.25 µM FMAL-actin and 0-20 µM TMR-cC0-C2 ±PKA (solid black line, unphosphorylated; dashed black line, PKA phosphorylated (#P)) and TMR-fskC1-C2 (solid blue line). (B) Expansion of 0-5 µM TMR-cC0-C2 ±PKA and TMR-fskC1-C2. The concentration of TMR-cC0-C2 and TMR-fskC1-C2 (0.5 µM) and phosphorylated TMR-cC0-C2 (1 µM) used in in the original screen and in testing of compound effects is indicated with arrows. Data are means ± SE. n=7-10 from N=2 separate actin and cC0-C2 or fskC1-C2 preparations. All values are the average ± SE.

### FLT of TMR-cC0-C2 biosensor

In addition to dramatically reducing the FLT-FRET between FMAL-F-actin and TMR-cC0-C2, the binding of the identified compounds to cC0-C2 changed the FLT of the directly excited acceptor probe, TMR on cC0-C2 (Figure S2). This suggested that using the same FLT-FRET samples containing labeled FMAL-F-actin and labeled TMR-cC0-C2 we could screen first for compounds that modulate cC0-C2 binding to actin by monitoring FMAL-F-actin FLT (excited at 473 nm), and then screen for compounds that bind to cC0-C2 by re-reading the plate and monitoring the FLT of TMR-cC0-C2 (excited at 532 nm).

### High-throughput screening of an FDA-approved library for compounds modulating cC0-C2 actin interactions and compounds that bind cC0-C2

Using the FMAL-F-actin plus TMR-cC0-C2 FRET biosensor, we performed HTS of the Selleck library that contains 2684 FDA-approved compounds. The entire screen was performed in duplicate with two different preparations of FMAL-F-actin and TMR-cC0-C2 samples. The Z’ factor (16), which validates the robustness of the FMAL-actin plus TMR-cC0-C2 FLT-FRET assay, was calculated as 0.61 for the first screen and 0.63 for the second using DMSO-only controls comparing over 600 wells with and without TMR-cC0-C2 (Table S1). The quality of the HTS screens were also validated by calculating the Z’ factor (0.62±0.01) using FMAL-F-actin plus TMR-cC0-C2 FLT-FRET without and with suramin (a tool compound identified in our earlier study), which is eliminating binding between actin and cC0-C2 (see experimental procedure).

Each screen monitored the effect of the compounds (1) on the FLT of FMAL-F-actin (donor only, D) at 473 nm, (2) on the FLT of FMAL-F-actin in the presence of TMR-cC0-C2 (donor-acceptor, DA) at 473 nm and (3) on the FLT of TMR-cC0-C2 (acceptor, A) in the presence of FMAL-actin at 532 nm (Figure 4). Two approaches to analyzing the data (see experimental procedures) yielded 60 hit compounds that either decreased (Table 1, Figure 4A) or increased (Table 2, Figure 4A) FLT-FRET between FMAL-F-actin and TMR-cC0-C2, and/or bound to TMR-cC0-C2 causing its FLT to decrease or increase (Table 3, Figure 4B). Compounds, which influence FRET between FMAL actin and cC0-C2 are potential effectors for actin-cMyBPC interactions, whereas compounds that impact FLT of TMR-cC0-C2 bind to cC0-C2 and likely affect its structure. These 60 compounds were selected to study concentration response effects. (Table S2)

**Figure 4.**
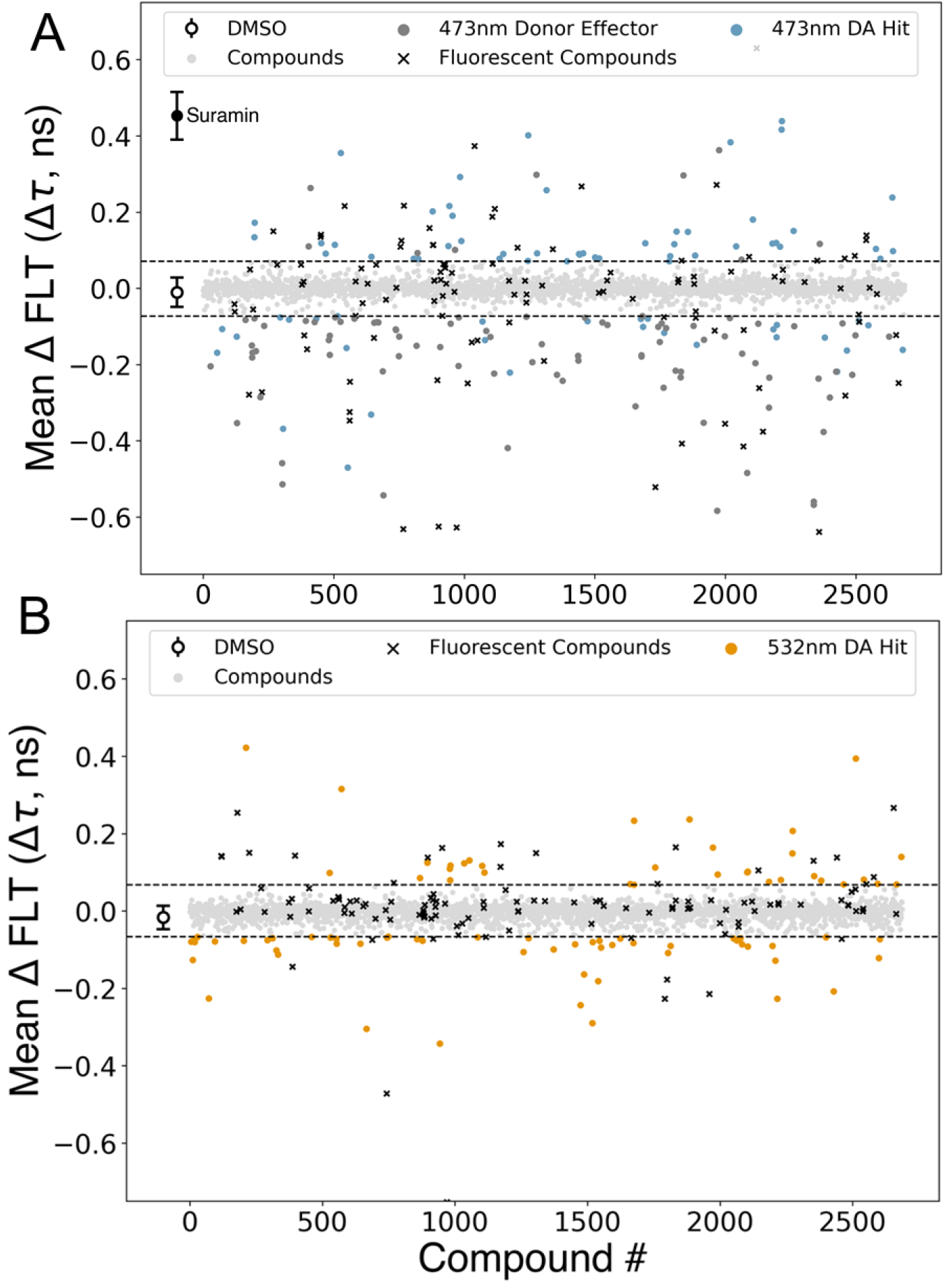
A representative screen of the Selleck library. Screen was done in duplicates with two different preparations of FMAL-F-actin and TMR-C0-C2 and reproducible hits are identified. Hits are identified as ±4 SD of the control (DMSO only) samples (A) Excitation at 473 nm. Reproducible donor only (dark gray dots) and DA (blue) hits at 473 nm. Donor only hits are excluded from the ‘hits’ list as they affect the lifetime of F-MAL-F-actin directly. Fluorescent compounds are shown in black and compounds that do not change FLT by >4 SD are shown in light grey. (B) Excitation at 532 nm. Reproducible DA hits are shown as orange dots.

**Table 1.**
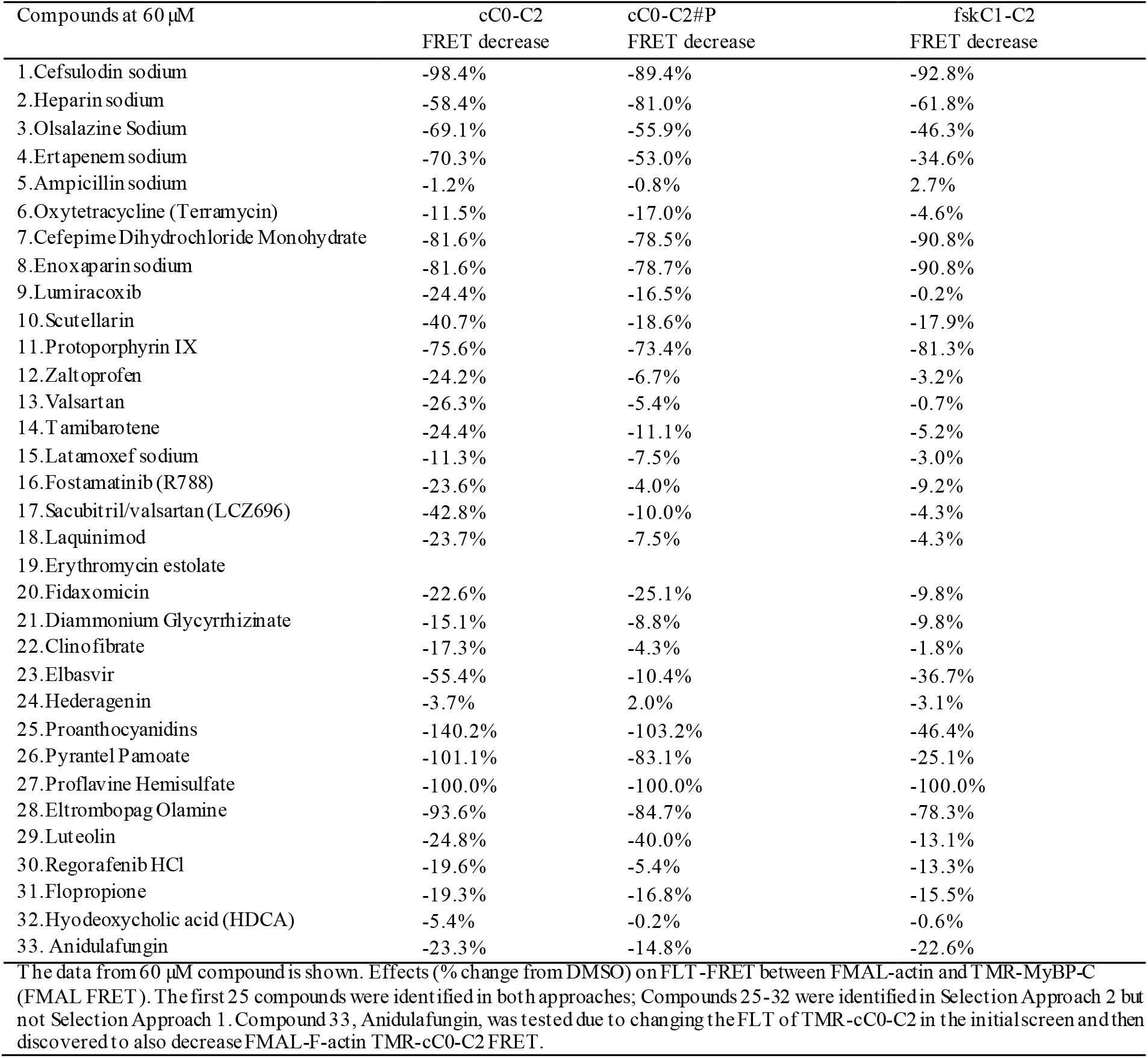
Compounds Decreasing FMAL-F-actin- TMR-cC0-C2 FLT-FRET (excited at 473 nm)

| Compounds at 60 μM | cC0-C2 | cC0-C2#P | fskC1-C2 |
| --- | --- | --- | --- |
|  | FRET decrease | FRET decrease | FRET decrease |
| 1.Cefsulodin sodium | -98.4% | -89.4% | -92.8% |
| 2.Heparin sodium | -58.4% | -81.0% | -61.8% |
| 3.Olsalazine Sodium | -69.1% | -55.9% | -46.3% |
| 4.Ertapenem sodium | -70.3% | -53.0% | -34.6% |
| 5.Ampicillin sodium | -1.2% | -0.8% | 2.7% |
| 6.Oxytetracycline (Terramycin) | -11.5% | -17.0% | -4.6% |
| 7.Cefepime Dihydrochloride Monohydrate | -81.6% | -78.5% | -90.8% |
| 8.Enoxaparin sodium | -81.6% | -78.7% | -90.8% |
| 9.Lumiracoxib | -24.4% | -16.5% | -0.2% |
| 10.Scutellarin | -40.7% | -18.6% | -17.9% |
| 11.Protoporphyrin IX | -75.6% | -73.4% | -81.3% |
| 12.Zaltoprofen | -24.2% | -6.7% | -3.2% |
| 13.Valsartan | -26.3% | -5.4% | -0.7% |
| 14.Tamibarotene | -24.4% | -11.1% | -5.2% |
| 15.Latamoxef sodium | -11.3% | -7.5% | -3.0% |
| 16.Fostatinib (R788) | -23.6% | -4.0% | -9.2% |
| 17.Sacubitril/valsartan (LCZ696) | -42.8% | -10.0% | -4.3% |
| 18.Laquinimod | -23.7% | -7.5% | -4.3% |
| 19.Erythromycin estolate |  |  |  |
| 20.Fidaxomicin | -22.6% | -25.1% | -9.8% |
| 21.Diammonium Glycyrrhizinate | -15.1% | -8.8% | -9.8% |
| 22.Clinofibrate | -17.3% | -4.3% | -1.8% |
| 23.Elbavir | -55.4% | -10.4% | -36.7% |
| 24.Hederagenin | -3.7% | 2.0% | -3.1% |
| 25.Proanthocyanidins | -140.2% | -103.2% | -46.4% |
| 26.Pyrantel Pamoate | -101.1% | -83.1% | -25.1% |
| 27.Proflavine Hemisulfate | -100.0% | -100.0% | -100.0% |
| 28.Eltrombopag Olamine | -93.6% | -84.7% | -78.3% |
| 29.Luteolin | -24.8% | -40.0% | -13.1% |
| 30.Regorafenib HCl | -19.6% | -5.4% | -13.3% |
| 31.Flopropione | -19.3% | -16.8% | -15.5% |
| 32.Hyodeoxycholic acid (HDCA) | -5.4% | -0.2% | -0.6% |
| 33. Anidulafungin | -23.3% | -14.8% | -22.6% |

The data from 60 μM compound is shown. Effects (% change from DMSO) on FLT-FRET between FMAL-actin and TMR-MyBP-C (FMAL FRET). The first 25 compounds were identified in both approaches; Compounds 25-32 were identified in Selection Approach 2 but not Selection Approach 1. Compound 33, Anidulafungin, was tested due to changing the FLT of TMR-cC0-C2 in the initial screen and then discovered to also decrease FMAL-F-actin TMR-cC0-C2 FRET.

**Table 2.** Compounds increasing FMAL-F-actin- TMR-C0-C2 FRET (excited at 473 nm)

|  |  |  |  |
| --- | --- | --- | --- |
| Compounds at 60 μM | cC0-C2 | cC0-C2#P | fskC1-C2 |
|  | FRET increase | FRET increase | FRET increase |
| 1.Clindamycin palmitate HCl | 54.1% | 69.6% | 34.8% |
| 2.Bedaquiline fumarate | 33.1% | 39.3% | 32.1% |
| 3.Tocofersolan | 26.1% | 1.9% | 6.3% |
| 4.Fenticonazole Nitrate | 5.9% | 29.5% | 1.5% |
| 5.Saikosaponin D | 22.3% | 0.5% | 5.6% |
| 6.Verteporfin | 47.0% | 8.3% | 2.1% |
| 7.Raloxifene | 112.8% | 81.4% | 75.6% |
| 8.Masitinib (AB1010) | 53.9% | 29.3% | 29.5% |
| 9.Doxazosin | 38.8% | 20.7% | 15.0% |
| 10.Telotristat Etiprate (LX 1606 Hippurate) | 15.8% | 27.6% | 14.4% |
| 11.Sanguinarine chloride | 9.1% | 12.4% | 3.6% |
| The data from 60 μM compound is shown. Effects (% change from DMSO) on FLT-FRET between FMAL-actin and TMR-MyBP-C (FMAL FRET). First 6 compounds were identified in both approaches; Compounds 7-11 were identified in Selection Approach 2 but not Selection Approach 1. | | | |

**Table 3.** Effect of the compounds on TMR-cC0-C2 FLT(excited at 532 nm)

| **Table 3. Effect of the compounds on TMR-cC0-C2 FLT(excited at 532 nm)** |  |  |  |
| --- | --- | --- | --- |
| Compounds at 60 μM | cC0-C2 | cC0-C2#P | fskC1-C2 |
|  | FLT | FLT | FLT |
| 1.Cefsulodin sodium | 9.1% | 8.1% | 11.5% |
| 2.Heparin sodium | 0.6% | 2.6% | 2.6% |
| 3.Olsalazine Sodium | -1.3% | -2.4% | -7.0% |
| 4.Ertapenem sodium | 10.2% | 8.0% | 12.2% |
| 5.Ampicillin sodium | -0.1% | -0.7% | -0.8% |
| 6.Oxytetracycline (Terramycin) | 2.4% | 4.9% | 2.1% |
| 7.Cefepime Dihydrochloride Monohydrate | 3.1% | 4.1% | 2.8% |
| 8.Enoxaparin sodium | 3.1% | 6.9% | 2.8% |
| 9.Lumiracoxib | 1.9% | 0.3% | -0.7% |
| 10.Scutellarin | -9.1% | -8.7% | -9.5% |
| 11.Protoporphyrin IX | -2.2% | 1.8% | -30.7% |
| 12.Zaltoprofen | 1.0% | 0.8% | 0.3% |
| 13.Valsartan | -0.3% | 0.5% | 0.0% |
| 14.Tamibarotene | 5.2% | 7.2% | 5.9% |
| 15.Latamoxef sodium | 0.8% | 3.6% | -1.5% |
| 16.Fostatinib (R788) | 2.3% | 0.7% | 0.4% |
| 17.Sacubitril/valsartan (LCZ696) | 0.8% | 1.2% | 2.6% |
| 18.Laquinimod | -1.6% | 0.1% | -0.8% |
| 19.Erythromycin estolate | -14.1% | -2.4% | -2.1% |
| 20.Fidaxomicin | -5.2% | -0.5% | -4.3% |
| 21.Diammonium Glycyrrhizinate | -0.3% | 1.6% | 2.1% |
| 22.Clinofibrate | 0.6% | 1.5% | 0.6% |
| 23.Elbavir | -1.0% | -3.7% | -3.5% |
| 24.Hederagenin | 2.5% | 3.6% | 2.3% |
| 25.Proanthocyanidins | -85.1% | -84.0% | -55.4% |
| 26.Pyrantel Pamoate | -3.7% | -1.4% | -7.9% |
| 27.Proflavine Hemisulfate | -1.0% | -1.3% | -1.1% |
| 28.Eltrombopag Olamine | -30.1% | -23.0% | -24.6% |
| 29.Luteolin | -15.3% | -14.6% | -13.0% |
| 30.Regorafenib HCl | 20.5% | 24.9% | 31.6% |
| 31.Flopropione | 0.6% | 0.1% | -0.7% |
| 32.Hyodeoxycholic acid (HDCA) | 1.5% | 1.5% | 0.6% |
| 33.Anidulafungin | 32.1% | 43.3% | 53.0% |
| 34.Clindamycin palmitate HCl | -12.7% | -10.2% | -1.2% |
| 35.Bedaquiline fumarate | -4.5% | -6.5% | 6.7% |
| 36.Tocofersolan | 3.3% | 3.2% | 2.8% |
| 37.Fenticonazole Nitrate | -4.7% | -6.0% | 0.4% |
| 38.Saikosaponin D | 5.4% | 6.3% | 4.2% |
| 39.Verteporfin* | 4.5% | 5.7% | 3.4% |
| 40.Raloxifene | -0.7% | -2.0% | -1.8% |
| 41.Masitinib (AB1010) | -5.0% | -2.5% | -2.1% |
| 42.Doxazosin | 8.1% | 6.5% | 5.4% |
| 43.Telotristat Etiprate (LX 1606 Hippurate) | 4.1% | 1.1% | 11.0% |
| 44.Sanguinarine chloride | 1.7% | 2.2% | 1.8% |

The data from 60 μM compound is shown. Effects (% change from DMSO) on the FLT of TMR-MyBP-C. Compounds 25, 26, 28, 29, 30, 41, 42, 43 were identified in Selection Approach 2 but not Selection Approach 1.

### Concentration response curves (CRC)

Sixty compounds identified as initial hits were further examined to determine the concentration response curves for seven conditions. (1) FMAL actin alone identified actin-binding compounds that alter the FLT of FMAL actin at higher concentrations. (2) FMAL actin plus TMR-cC0-C2, focusing on the FLT of FMAL as in the initial screen, monitored compound effects on actin-cC0-C2 interactions. (3) FMAL actin with PKA-phosphorylated TMR-cC0-C2, focusing on the FLT of FMAL, monitored compound effects on interaction of actin with phosphorylated cC0-C2. (4) FMAL actin plus TMR-fskC1-C2, focusing on the FLT of FMAL, monitored compound effects on actin-fskC1-C2 binding. (5) TMR-cC0-C2, focusing on the FLT of TMR, monitored cC0-C2-binding compounds. (6) PKA phosphorylated TMR-cC0-C2 focusing on the FLT of TMR monitored phosphorylated cC0-C2-binding compounds. (7) TMR-fskC1-C2 focusing on the FLT of TMR, monitored fskC1-C2-binding compounds. CRC for representative compounds of each category are shown in Figure 5. Out of these 60 compounds, 22 either showed very little effect on CRC curves or increased the donor only lifetime at the higher compound’s concentration. The effects of remaining 38 compounds that showed reasonable CRC responses, alongside the 22 other compounds, are summarized in Tables 1-3 with their FRET/FLT changes at 60µM.

**Figure 5.**
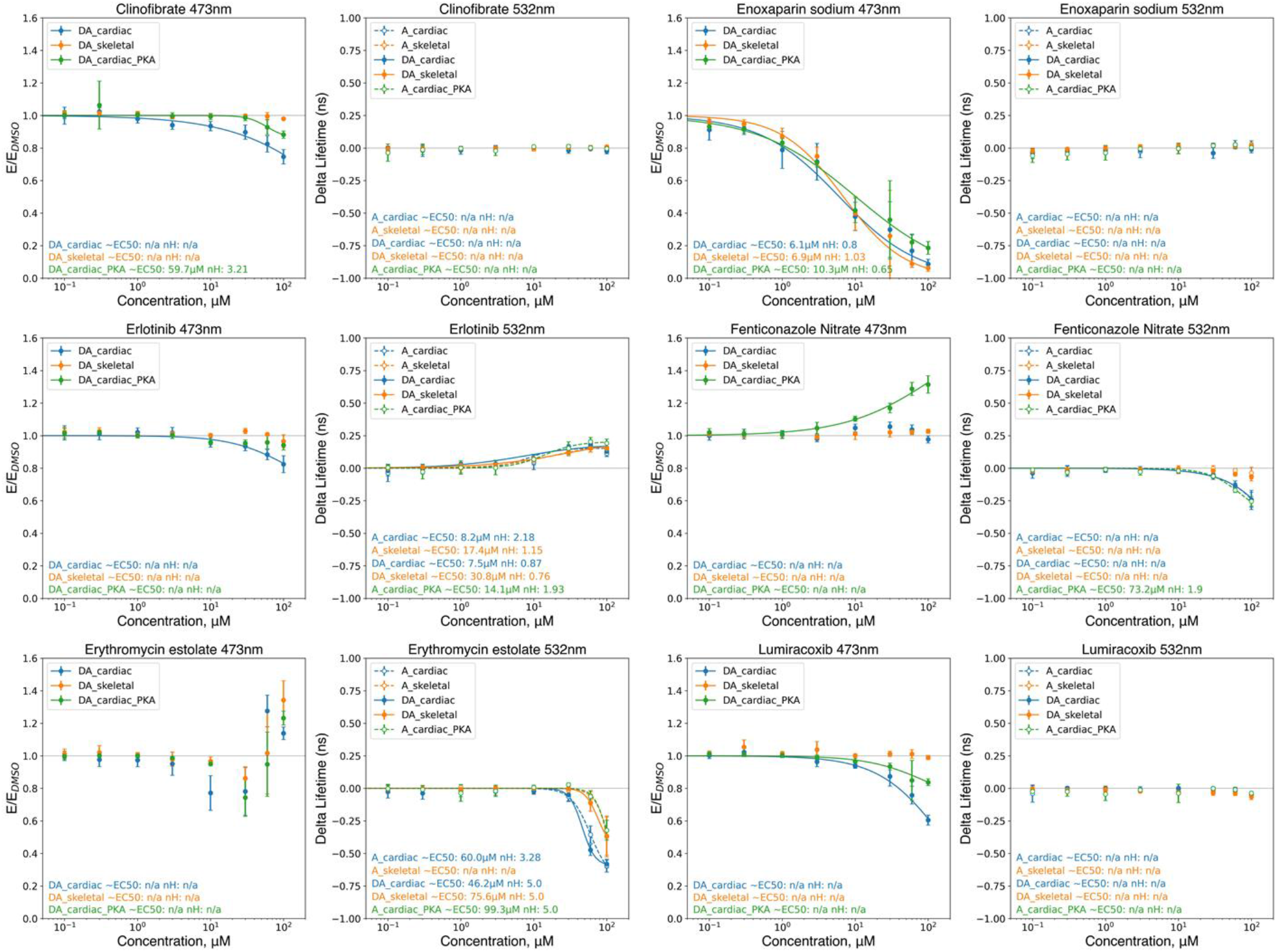
FLT-FRET Concentration Response Curves (CRC) of the selected hits, excited at 473 nm or 532 nm. Clinofibrate, Enoxaparin sodium, Erlotinib, Lumiracoxib decrease the FLT-FRET between FMAL-F-actin and TMR-cC0-C2.Fenticonazole nitrate increases FLT-FRET between FMAL-F-actin and TMR-cC0-C2#P but does not have any effect on cC0-C2 or fSkC1-C2. Erthromycin estolate has a biphasic effect on FRET between FMAL-F-actin and TMR-cC0-C2/TMR-cC0-C2#P/TMR-fskC1-C2. Erlotinib increases TMR-C0-C2 FLT for all samples, and Erthromycin estolate decrease them. Fenticonazole nitrate slightly decreases FLT of TMR-cC0-C2 and TMR-cC0-C2#P but does not affect TMR-fskC1-C2. Data are collected from three independent preparation of actin and TMR-cC0-C2 and TMR-fskC1-C2. n=9, N=3. The 60 M results are summarized in Table 1, 2 and 3. Error is in SD.

33 initial hits that decreased FMAL-actin TMR-cC0-C2 FRET in the initial screen again showed reduced FLT-FRET (Table 1). Ampicillin and Hederagenin were exceptions. Lumiracoxib, Valsartin, Erythromycin estolate, Clinofibrate, and Hyodeoxycholic acid (HDCA) were cardiac specific, showing no effect on FMAL-F-actin TMR fskC1-C2 FLT-FRET. Seven other compounds, Zaltoprofen, Tamibarotene, Latamoxef sodium, Sacubitril/Valsartan (LCZ696), Laquinimod, Proanthocyanidins, and Pyrantel Pamoate showed a marked preference for reducing FLT-FRET effects, with cC0-C2 reductions being 3-10 times that of fskC1-C2. Hederagenin and HDCA, appeared to reduce (slightly) actin binding of unphosphorylated cC0-C2, or fskC1-C2 but not phosphorylated cC0-C2. Seven compounds showed a 4-7 times greater reduction in cC0-C2 than phosphorylated cC0-C2. These compounds were Zaltoprofen, Valsartin, Fostamatinib (R788), Sacubitril/valsartan (LCZ696). Finally, 23 of the 33 compounds in this group affected the FLT of TMR on cC0-C2 and/or fsk, indicative to binding to these proteins (Table S4). Twelve of the compounds altered the FLT of FMAL-actin by 1.4% or more and likely bind to actin.

The 11 compounds that were identified as increasing FMAL-actin FRET in the initial screen again showed increased FRET (Table 2). All 11 altered the FLT of TMR on cC0-C2 and/or fskC1-C2 indicating that they bind to MyBP-C. Fenticonozole Nitrate showed the strongest (5 times) preference for effecting binding of the phosphorylated over unphosphorylated cC0-C2. Tocofersolan and Saikosaponin D affected the unphosphorylated cC0-C2 and fskC1-C2 binding to actin but not the phosphorylated cC0-C2 despite binding to it (see changes in TMR FLT). Clindamycin palmitate HCl, Bedaquiline fumarate, Verteporfin and Sanguinarine chloride increased FLT-FRET levels for all 3 MyBP-C proteins (Table S3). Raloxifene, Masitinib (AB1010), Doxazosin, and Telotristat Etiprate (LX 1606 Hippurate) also increased FLT-FRET, but these displayed effects on FMAL-F-actin alone and were removed from further consideration.

8 compounds that reduced the FLT of TMR-cC0-C2 in the initial screen only one, Scutellarin, reduced it when combined with all three MyBP-C fragments in the test for CRC (Table 3). This may indicate that the reduction in FLT of TMR-cC0-C2 by 8-hydroxyquinoline and Benzonate was dependent on the presence of actin in the initial screen. Five of these compounds were removed from consideration due to effects on FMAL-F-actin alone.

The 20 compounds that increased the FLT of TMR-cC0-C2 (Table 3) all retested in CRC tests as doing the same. Four of the compounds altered the FLT of FMAL-actin by 2.5% or more and were removed from consideration. The remaining compounds displayed no large differences between effects on TMR-cC0-C2, phosphorylated TMR-cC0-C2, or TMR-fskC1-C2.

Thus, we identified different groups of effectors that modulate either the structural complex of actin-cC0-C2 and/or bind to cC0-C2.

### Cardiac and Skeletal ATPase assays

Compounds that showed significant CRC effects on the structural assays were further tested for their effects on muscle function. Compound effects were examined on the Ca^2+-^dependent ATPase activity of bovine cardiac (left ventricle) myofibrils (BvCMF) and rabbit fast skeletal (psoas) myofibril (RSkMF) in a moderate-throughput manner, varying the free calcium concentration from pCa 8 (relaxation) to pCa 4 (full activation). The effects of 21 current hits and two previously identified compounds (NF023 and Suramin) were tested for their effects on the myosin ATPase activity in BvCMF and RSkMF. The pCa_50_ values of BvCMFs and RSkMF are 5.76 ± 0.02 and 6.06 ± 0.01 respectively, consistent with previous reports (17). Suramin (from the previous study) had the most prominent effects on both BvCMF and RSkMF ATPase. Compounds, identified in this current screen such as Pneumocandin B0, and Enoxaparin sodium, showed marked differences in ATPase levels and/or Ca^2+^ sensitivity in BvCMF and RSkMF. Pranlukast and Erlotinib also showed differences. ATPase results are summarized in Table 4 and the representative compounds, which show distinct differences between the skeletal and cardiac myofibrils are shown in Figure 6.

**Table 4:** Summary of myofibril ATPase activity.

| Compound | BvCMF |  | RSkMF |  |
| --- | --- | --- | --- | --- |
|  | Avg pCa50 | SD | Avg pCa50 | SD |
| **DMSO** | **5.782** | **0.021** | **6.055** | **0.013** |
| Cefsulodin sodium | 5.701 | 0.008 | 6.026 | 0.280 |
| Pranlukast | 5.703 | 0.025 | 6.104 | 0.078 |
| Ertapenem sodium | 5.731 | 0.030 | 5.969 | 0.015 |
| Tamibarotene | 5.753 | 0.032 | 6.102 | 0.005 |
| Valsartan | 5.795 | 0.053 | 6.062 | 0.036 |
| Clinofibrate | 5.818 | 0.051 | 6.055 | 0.022 |
| Tocofersolan | 5.765 | 0.052 | 6.100 | 0.010 |
| Erlotinib | 5.677 | 0.134 | 6.017 | 0.035 |
| Cefepime Dihydrochloride Monohydrate | 5.772 | 0.011 | 6.085 | 0.021 |
| Zaltoprofen | 5.810 | 0.013 | 6.086 | 0.002 |
| Laquinimod | 5.793 | 0.012 | 6.067 | 0.056 |
| Fenticonazole Nitrate | 5.812 | 0.010 | 6.029 | 0.011 |
| Fostamatinib (R788) | 5.789 | 0.068 | 6.084 | 0.015 |
| Heparin sodium* | 5.707 | 0.071 | 6.062 | 0.036 |
| a-Hederin | 5.820 | 0.040 | 6.044 | 0.001 |
| Pneumocandin B0 | 5.927 | 0.127 | 6.118 | 0.016 |
| Saikosaponin D | 5.764 | 0.014 | 6.013 | 0.024 |
| Troxerutin | 5.794 | 0.024 | 6.058 | 0.018 |
| Triclocarban | 5.790 | 0.030 | 6.058 | 0.032 |
| Chloroquine Phosphate | 5.781 | 0.043 | 6.050 | 0.029 |
| NF023 | 5.670 | 0.012 | n.d. | n.d. |
| Suramin | 4.712 | 0.001 | 5.776 | 0.082 |
| Verteporfin | 5.858 | 0.203 | n.d. | n.d. |
| Enoxaparin sodium | 5.560 | 0.026 | 6.058 | 0.006 |

Compound concentration was 50 μM except \* (10 μM). All values are the average ± SD. N=3 for BvCMF (n=6 for each $\text{Ca}^{2+}$ concentrations) and N=2 for RSkMF (n=4)

**Figure 6.**
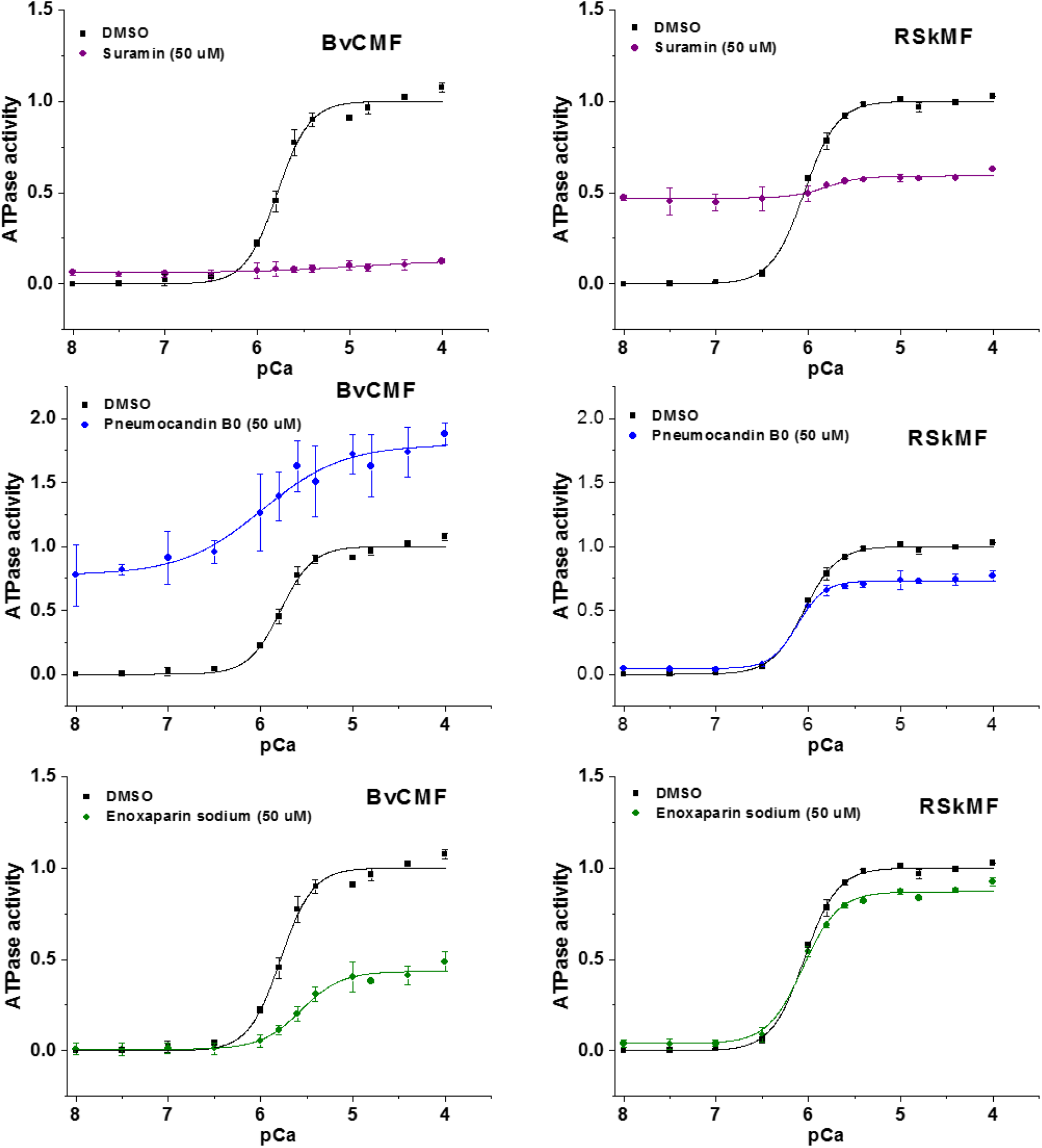
Myofibril ATPase assay. ATPase activity of bovine cardiac myofibrils (BvCMF) and rabbit skeletal (RSkMF) were measured in the presence of 1% DMSO and 50 µM compound across a 12-point Ca^2+^ gradient, pCa 4-8, in a 96-well–plate. Error bars are SD (N = 3, n=6).

## Discussion

We report a novel approach to discover two distinct classes of compounds that affect the binding of N-terminal cC0-C2 fragment of cMyBP-C to actin and myosin, with the ultimate goal of developing improved therapies for HF patients. We have developed two HTS screens that successfully identified a number of compounds capable of influencing muscle regulation by binding to MyBP-C.

The principal goal of this study is to identify novel small molecule effectors with therapeutic potential for disorders associated with mutations in sarcomeric proteins, including HF. Small-molecule effectors designed to target striated muscle proteins, such as myosin and troponin for the treatment of disease are showing assurance in preclinical and clinical trials (12,18). In our work, we focus on modulation of cMyBP-C interactions with both actin and myosin. This approach is novel as targeting cMyBP-C offers many of the same advantages for modulating heart function as targeting myosin directly, with the additional benefit of being a cardiac-specific accessory protein. As such, we expect drugs binding to cMyBP-C to result in fewer side effects, medical complications, or long-term effects.

Compounds identified from the Selleck library that altered the interaction between F-actin and cC0-C2 or bound to cC0-C2 were detected with high precision. The optimized HTS assay using FMAL-F-actin and TMR-cC0-C2 showed a Z’ value of 0.6 ± 0.1 in 1536 well format, which is considered excellent in the field of HTS assay development. We selected 60 hits from the primary FLT-FRET and FLT-TMR assays for an overall hit rate of 2%. Thirty-two (1.1%) reduced and 8 (0.3%) increased FLT-FRET between F-actin and cC0-C2. Twenty-eight (1.0%) displayed cC0-C2 binding.

CRC analysis of compound effects on actin alone removed 17 of the initial 60 selected hit compounds as they showed binding to actin alone. Three compounds did not show activity upon retesting in the CRCs. These 22 were removed from further consideration, leaving 38 potential hits.

The use of 6 additional CRCs performed on the remaining 41 hits allowed the characterization of them based on the specificity of their effects on cardiac versus skeletal MyBP-C and on unphosphorylated versus phosphorylated MyBP-C. For actin binding effects, five compounds showed isoform specificity where decreased binding was observed for cC0-C2 but not fskC1-C2. Seven compounds showed a strong (3-10x) preference for effects on cC0-C2 versus fskC1-C2 actin binding. Nine compounds showed more (4-7x) reduction in effects on actin binding of unphosphorylated than phosphorylated cC0-C2.

The interpretation of FLT-FRET and FLT-TMR values are very different. For FLT-FRET, the magnitude and direction of FLT-FRET changes correlate with cC0-C2 or fskC1-C2 interactions with actin. Increased FLT-FRET indicates increased binding and decreased FLT-FRET indicates decreased binding. For FLT-TMR, changes of TMR on cC0-C2 or fskC1-C2 as compounds bind to these proteins means only that the compound is binding and somehow altering the lifetime of TMR. This value can be positive or negative and large or small such that a 10x difference in FLT does not indicate a 10x difference in binding levels. Also, lack of a change of FLT does not mean lack of binding. Compounds can bind to cC0-C2 or fskC1-C2 without affecting the lifetime of TMR. The same is true for the magnitude of FLT changes of FMAL on actin in the absence of cC0-C2 or fskC1-C2.

Effects of 21 hits from this screen and suramin and NF023, from the previous study (13) were tested for their ability to modulate Ca^2+^ stimulated myosin ATPase in cardiac as well as in skeletal myofibril. Thirteen compounds initially selected for their modulation of C0-C2 actin interactions (Tables 1 and 2) and 11 originally selected for binding to cC0-C2 in the FLT-TMR assay (Table 3) were tested. Four of the compounds fell into both categories. Six of the 21 tested compounds affected ATPase rates at either low or high calcium or influenced the pCa50 values for the BvCMF. Suramin and NF023, identified previously, had strong effects on cardiac myofibril ATPase activity. These compounds also modulated the RSkMF activity; however, there is a difference in the degree of inhibition or activation or shift in Ca^2+^ sensitivity betweenthe two muscle types.

For those compounds that altered the myofibril ATPase activity, there is no clear correlation with the effects on cC0-C2-F-actin binding as reported by FLT-FRET. This is not surprising, as myofibril consists of several proteins as well as full length MyBP-C. In this final group of 6 compounds, heparin sodium, Enoxaparin sodium, Pranlukast, show large decreases in cC0-C2 interactions with actin, two, Pneumocandin BO and α-hederin, show increases in cC0-C2 and fskC1-C2 but not phosphorylated cC0-C2, and one, erlotinib, shows only modest decreases in cC0-C2 interactions with actin. That identified compounds either increase or decrease FRET, indicates that the compounds can either enhance or reduce the binding between actin and cC0-C2. We speculate two possible modes of action for these compounds: 1. binding to cC0-C2 either increases or decreases the binding of MyBP-C to actin, altering thin filament conformation resulting in cooperative effects on the Ca^2+^ sensitivity and 2. binding to cC0-C2 could also alter its myosin interactions thereby modulating myosin’s activation state under different conditions.

Much remains to be learned about the effects of these compounds on cardiac muscle contractility, but the approach used in this work clearly indicates that the primary and secondary assays are suitable for screening thousands of compounds to identify cMyBP-C binding hits capable of altering its function. We expect that these compounds will be valuable tools for understanding drug targeting and mechanism of action (MOA) of cMyBP-C to reduce contractility in HCM or enhance it in DCM. A key bottleneck for progressing to clinical trials is the development of drugs that bind with high affinity with the desired target. Our compounds affect muscle regulation at micromolar concentrations, indicating moderate binding affinity, which can be optimized in the future through medicinal chemistry. Modulation of structures of these compounds through medicinal chemistry will also provide valuable information for predicting efficacy. The compounds identified in this screen are already FDA approved and in use as medications for several diseases and some have significant side effects. An effective drug must achieve a balance between specific therapeutic benefit and undesired side effects. Future screening of larger libraries, with the current assays developed in this study, will also help identify variable scaffolds that can safely work on a particular muscle type.

### Experimental Procedures

#### Actin preparations and labeling

Actin was prepared from rabbit skeletal muscle by extracting acetone powder in cold water, as described previously (13). 50 μM of G-actin in 20 mM Tris pH 7.5, 0.2 mM CaCl_2_, 0.2 mM ATP was polymerized by the addition of 3M KCl (to a final concentration of 100 mM) and 0.5 M MgCl_2_ (to a final concentration of 2 mM), followed by incubation at 23 ºC for 1 hour. Labeling with FMAL was done at a final FMAL concentration of 1 mM for 5 hours at 23 ºC and then overnight at 4 ºC. Labeling was stopped by the addition of a five-fold molar excess of DTT. Unincorporated dye was removed by cycling the actin through F-actin and G-actin states as described in (15). To avoid FLT-FRET between FMAL on the neighboring Cys-374 residues of actin monomers in F-actin, unlabeled G-actin was mixed with the FMAL-actin to achieve ∼10% FMAL-actin prior to the final actin polymerization. Finally, fluorescent-labeled F-actin was stabilized by the addition of equimolar phalloidin.

For all assays F-actin was resuspended in and/or dialyzed against MOPS-actin binding buffer, M-ABB (100 mM KCl, 10 mM MOPS pH 6.8, 2 mM MgCl_2_, 0.2 mM CaCl_2_, 0.2 mM ATP, 1 mM DTT).

#### Recombinant human preparations and labeling. cMyBP-C fragment

pET45b vectors encoding *E. coli* optimized codons for the cC0-C2 portion of human cMyBP-C with N-terminal 6x His tag and TEV protease cleavage site were obtained from GenScript (Piscataway, NJ). For FLT-FRET binding assays, we mutated cC0-C2 so that it contained a single cysteine at position 249, a surface-exposed residue in the C1 domain. To achieve this, 4 endogenous cysteines in cC0-C2 were mutated to amino acids found in other MyBP-C proteins leaving only one cysteine at position 249 (19). Protein production in *E. coli* BL21DE3-competent cells (New England Bio Labs, Ipswich, MA) and purification of cC0-C2 protein using His60 Ni Superflow resin was done as described (20). cC0-C2 (with His-tag removed by TEV protease digestion) was further purified using size-exclusion chromatography to achieve >90% intact cC0-C2 as described (21) and then concentrated, dialyzed to 50/50 buffer (50 mM NaCl and 50 mM Tris, pH 7.5) and stored at 4 °C.

For FLT-FRET experiments, cC0-C2^Cys249^ was labeled with tetramethylrhodamine (TMR) in 50/50 buffer. cC0-C2^Cys249^ (50 μM) was first treated with the reducing agent TCEP (200 μM) for 30 minutes at 23 ºC, and then TMR was added (from a 20 mM stock in DMF) to a final concentration of 200 μM. Labeling was done for 1 hour at 23 ºC and terminated by the addition DTT (to 1 mM). Unincorporated dye was removed by extensive dialysis against M-ABB buffer. The degree of labeling was kept at 0.8 dye/cC0-C2 as measured by UV-vis absorbance to avoid aggregation.

The N-terminal domains from fskMyBP-C, C1-C2 (skeletal MyBP-C does not contain the C0 domain present in cMyBP-C) were similarly expressed in bacteria and labeled with TMR.

#### In vitro phosphorylation of cMyBP-C

cC0-C2 was treated with 7.5 ng PKA/μg cC0-C2 at 30 ºC for 30 min, as described in (15,21).

#### Fluorescence data acquisition

Fluorescence lifetime (FLT) measurements were acquired using a high-throughput FLT plate reader (FLTPR; Fluorescence Innovations, Inc., Minneapolis, MN) (14,20), provided by Photonic Pharma LLC (Minneapolis, MN). For FLT-FRET experiments, donor FMAL actin and the donor+acceptor sample (FMAL actin+ TMRcC0-C2) were excited with a 473-nm microchip laser (Bright Solutions, Cura Carpignano, Italy) and emission was filtered with 488-nm long pass and 517/20-nm bandpass filters (Semrock, Rochester, NY). For FLT-TMR experiments, DA and A samples were excited at 532 nm. The observed waveforms were analyzed as described previously (15,22).

Plates were also scanned using the spectral unmixing plate reader (SUPR; Fluorescence Innovations, Inc., Minneapolis, MN) to exclude fluorescent compounds, which can present as false positives in the FLT-FRET measurement (14,23)

#### FMAL-Actin-TMR-MyBP-C FLT-FRET binding assays

FLT-FRET binding assays were performed as described (13) with the modification being that 0.25 μM F-Actin labeled (to 10%) on Cys-374 with FMAL (donor) was incubated with cC0-C2 Cys^249^, fskC1-C2 or sskC1-C2 labeled to 70-90% with TMR (acceptor) (for donor-acceptor (DA) data). FLTs of FMAL-actin were determined for D and DA. FLT-FRET efficiency (1 – (τ_DA_ / τ_D_)) was determined.

The binding curve (Figure 3) was generated from data from 2 separate preparations of actin and cC0-C2^Cys249^. Testing of compound effects by FLT-FRET was similarly done at 1 µM FMAL-actin with unlabeled C0-C2Cys225 (D) and TMR-C0-C2Cys225 (DA), both at 2 µM.

#### Selleck library screen

2684 Selleck compound (Cat# L1300-Z394273, mostly FDA-approved drugs) were received in 96-well plates and reformatted into 1536-well flat, black-bottom polypropylene plates (Greiner Bio-One). In total, 50 nl of each solution was dispensed in DMSO using an automated Echo 550 acoustic liquid dispenser (Labcyte). Compounds were formatted into the assay plates, at a final concentration of 30 μM, with the first two and last two columns loaded with DMSO only (compound-free controls). These assay plates were then heat-sealed using a PlateLoc Thermal Microplate Sealer (Agilent Technologies) and stored at –20⁰C. Before screening, compound plates were equilibrated to room temperature (25⁰C). 0.25 μM F-MAL actin without or with 0.5 µM TMR-cC0-C2 was dispensed into 1536-well assay plates containing the compounds by a Multidrop Combi Reagent Dispenser (Thermo Fisher Scientific. Plates were incubated at room temperature for 60 min before recording the data with the FLTPR. Screens were performed in duplicates with independent preparations of actin and cC0-C2.

#### FLT data analysis

Following data acquisition, time-resolved FLT waveforms observed for each well were convolved with the instrument response function (IRF) to determine the FLT (τ) (Eq. 1) by fitting to a single-exponential decay (15,22,24).

The decay of the excited state of the fluorescent dye attached to actin at Cys-374 to the ground state is:

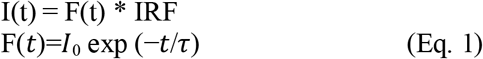

Where I is the measured waveform, *I*_*0*_ is the fluorescence intensity upon peak excitation (*t* = 0) and τ is the FLT (t = τ when *I* decays to 1/e or ∼37% of *I0*).

FLT assay quality was determined using two approaches. The first approach was the same as in our previous screen (13) This used controls (DMSO-only samples) on each plate as indexed by the *Z’* factor.

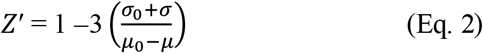

where σ_0_ and σ are the standard deviations (SD) of the controls τ_0_ and τ, respectively, and μ_0_ and μ are the means of the controls τ_0_ and τ, respectively.

The second Z’ factor calculation makes use of a tool compound as we have done previously (25). Here, assay quality was determined based on FRET assay samples in wells pre-loaded with control (DMSO) and tested tool compound (suramin), as indexed by the Z’ factor:

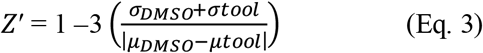

where σ_DMSO_ and σ_Tool_ are the SDs of the DMSO τ_DA_ and tool compound τ_DA_, respectively; µ_DMSO_ and µ_Tool_ are the means of the DMSO τ_DA_ and tool compound τ_DA_, respectively.

A Z’ factor value of 0.5 to 1 indicates excellent assay quality (15,16,26). Both approaches resulted in a Z’ of >0.6. A compound was considered a Hit if it changed τ_DA_ by > 4 SD relative to that of control τ_DA_ that were exposed to 0.3% DMSO.

#### HTS data analysis

Identification of the preliminary hits was done by two independent approaches.

Approach 1: First the compounds interacting with actin alone were removed from consideration. Those compounds in each plate that changed the FLT of FMAL on actin in the absence of cC0-C2 by more than 3 SD of the average FLT in wells (224 in plate 1 and 2, 176 in plate 3) that contain DMSO. Next, the FRET efficiency was calculated using the FLT of FMAL-F-actin (D) and FMAL-F-actin in the presence of TMR-cC0-C2 (DA) in the presence of individual compounds as in Equation 4. Standard Z-scores were determined for compounds in each plate using the average and SD of FLT-FRET for wells on the same plate that contain DMSO. Compounds that had Z-scores greater than or equal to 3 or less than or equal to -3 in both the first and screens were inspected to remove those known or found to be fluorescent (described in Approach 2) or aggregators. The resulting compounds were considered as actin-cC0-C2 interaction hits. This resulted in 34 that decreased FMAL-actin-TMR-cC0-C2 FRET and 9 that increased FMAL-actin-TMR-cC0-C2 FRET.

Hits that bind to TMR-cC0-C2 and change the FLT of TMR were similarly identified. The TMR FLTs of those compounds that remained, after removal of actin-interactors, were used to calculate Z-scores. Z-scores were determined for compounds in each plate using the average and SD of the TMR FLT for wells on the same plate that contain DMSO. Compounds that had Z-scores greater than or equal to 3 or less than or equal to -3 in both the first and screens were inspected to remove those known to be fluorescent or aggregators. The resulting compounds were considered cC0-C2 interaction hits. This resulted in 19 cC0-C2-binding preliminary hits. Three reduced and 16 increased the FLT of TMR-cC0-C2.

Approach 2: After fitting waveforms with a single exponential decay to quantify donor FLT, the change in FLT (Δτ) was computed by performing a moving median subtraction in the order the plate was scanned with a window size of half a plate row (24 columns). The reasons for this are twofold. Firstly, plate gradients are often observed due to heating of the digitizer during acquisition, and performing Δτ computations with DMSO controls alone can sometimes result in artifacts as half of the DMSO wells are on the edge of the plate, which occasionally exhibit artifacts due to processes needed for the preparation of the compound library being tested. As most compounds are likely to be non-hits, and therefore DMSO like, computation of a moving median is an effective alternative to solving both gradient issues and edge-effect distortion of the primary metric for hit selection, Δτ. A complementary metric, FRET efficiency (E), was also computed as the fractional decrease of donor FLT in the absence and in the presence of acceptor as in Equation 4:

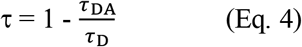

To account for the necessity to correct for plate gradients in the computation of E, the median τ_DMSO_ prior to application of the moving median correction was added to each well’s Δτ value to recover a rescaled value of τ. As the screen was run in duplicate, the mean values of τ and Δτ were used in the computation of metrics for hit selection. Ηit were selected by computing the robust *z*-score on a plate-by-plate basis, with a hit threshold set at ±4. The robust z-score, where the median (*M*) and median absolute deviation (*MAD*) are used in place of the mean and standard deviation (Equation 5), was used to best capture the most hits, as the standard z-score is more subject to strong outliers.

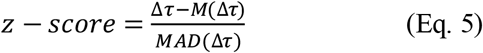

Two potential sources of false positives are compounds that either affect the donor only FLT directly or exhibit fluorescence under the conditions of the screen. Hits that exceeded the z-score hit criteria in the donor-only screen were excluded from consideration. To remove compounds with confounding fluorescence property, a similarity index (SI, specifically the cosine distance, Εq. Z) was computed using data from the SUPR instrument by comparing a region (500-540nm) of the donor only spectrum (*I*^(a)^) for each well to that of the plate-wide average DMSO spectrum (*I*^(b)^) in the same wavelength band (23). Compounds that exceeded an SI robust z-score of 4 were deemed likely fluorescent compounds and removed from consideration.

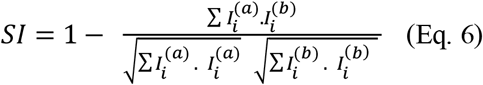

#### Concentration-response (CRC) assay

The primary Hits were dissolved in DMSO to make a 10 mM stock solution, which was serially diluted in in 96-well mother plates. Hits were screened at eight concentrations (0.5 to 100 μM). Compounds (1 μl) were transferred fromthe mother plates into 384-well plates using a Mosquito HV liquid handler (TTP Labtech Ltd., Hertfordshire, UK). The same procedure of dispensing as for the pilot screening was applied in the concentration-response assays. Concentration dependence of the FLT-FRET or FLT-TMR change was fitted using the Hill equation (27):

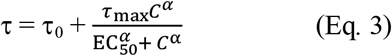

where τ and τ_0_ are FLT in the presence and in the absence of the compound, τ_max_ is the maximum effect, *C* is the compound concentration, EC_50_ is the compound concentration for which 50% of maximum effect is obtained, and α is the Hill coefficient of sigmoidicity.

#### Myofibril ATPase kinetics

Rabbit psoas and bovine cardiac myofibrils were prepared as described in our previous study (17). An enzyme-coupled, NADH-linked ATPase assay was used to measure skeletal and cardiac myofibril ATPase activity in 96-well microplates at 23⁰C. Ionic strength was maintained below 0.2 M, so myofilaments remain insoluble and stable (28). To achieve optimal enzymatic activities, buffers for skeletal and cardiac myofibrils differed slightly (29,30): skeletal in 50 mM MOPS, 100 mM KCl, 5 mM MgCl_2_, 1 mM EGTA, pH 7.0; cardiac in 20 mM MOPS, 35 mM NaCl, 5 mM MgCl_2_, 1 mM EGTA, pH 7.0. Each well in the microplate contained an assay mix of 0.84 mM phosphoenolpyruvate, 0.17 mM NADH, 10 U/vol pyruvate kinase, 20 U/vol lactate dehydrogenase. The concentration of free Ca^2+^ was controlled by EGTA buffering (31). Myofibrils were dispensed in microplates and were incubated with assay mix plus compound of interest for 20 minutes at 23°C. The concentration of myofibrils used were 0.01 mg/ml for skeletal and 0.05 mg/ml for cardiac. The assay was started upon the addition of ATP, at a final concentration of 2.1 mM (total volume to 200 µL), and absorbance was recorded at 340 nm in a SpectraMax Plus microplate spectrophotometer from Molecular Devices (Sunnyvale, CA). The steady-state myofibrillar ATPase rate was measured from NADH oxidation, measured as the rate of decrease in absorption at 340 nm for 30 minutes, with absorption data collected every 15 seconds. All data and statistical analysis of steady-state kinetics were conducted with the OriginPro program. The results were fitted with the Hill equation:

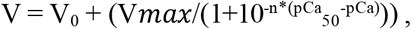

where V is the ATPase rate and n is the Hill coefficient.

ATPase activity is reported on the µmol ATP/mg protein/min scale. Average data are presented as mean ± SD. Sample means are derived from three different myofibrils preps (N=3), each done in triplicate (n=9).

The effects of compounds on ATPase activity of myofibrils were determined in the presence of 50 μM compound, across a 12-points pCa range from 4.0 (activation) to 8.0 (relaxation). The calcium sensitivity (pCa_50_) was determined by fitting the curves to the Hill equation. In all experiments, a 1% DMSO-only control was included.

#### Statistics

Average data are provided as mean ± standard error (SE) and each experiment was done with 3 separate protein preparations (N=3) and performed in triplicate (n=9) except RSkMF ATPase where N=2 (n=6).

## Data Availability

All data discussed are presented within the article.

## Author Contributions

B.A.C., D.D.T., T.A.B., and P.G. designed the study and wrote the paper. T.A.B. and V.C.L., purified recombinant proteins and DNA. T.A.B., V.C.L., P.G., and A.L.C. undertook purification of actin filaments for FLT assays. T.A.B, A.L.C. and P.G. conducted FLT assays, T.A.B optimized C0-C2^cys249^ labeling. A.L.C. prepared compound plates. T.A.B., B.A.C., P.G.., A.R.T., and D.D.T. analyzed FLT data and assisted with preparation of the figures and tables. V.C.L. and A.R.T. assisted with critical evaluation of the results and edited the manuscript. All authors critically evaluated and approved the final version of the manuscript.

## FOOTNOTES

## Acknowledgements

This work was supported by SBIR grant R43 HL162329 to B.A.C., D.T.T, and J.J.T. We thank Samantha Yuen for technical assistance with the FLT plate reader. We thank Bengt Svensson for assistant with graphics.

## Abbreviations

DCM: dilated cardiomyopathy
HCM: hypertrophic cardiomyopathy
cMyBP-C: cardiac myosin-binding protein C
P/A: proline/alanine-rich linker between domains C0 and C1
M: M-domain, phosphorylatable linker between C1 and C2
cC0-C2: N-terminal fragment of cMyBP-C comprised of C0-P/A-C1-M-C2 domains and linkers
HTS: high-throughput screen
FLT: fluorescence lifetime
FLTPR: FLT plate reader
FRET: fluorescence resonance energy transfer
FLT-FRET: FLT-detected FRET
DO: donor only
DA: donor plus acceptor
LOPAC: library of pharmacologically active compounds
F-actin: filamentous actin
G-actin: globular actin
ITC: isothermal titration calorimetry
TPA: transient phosphorescence anisotropy
K_d_: dissociation constant
B_max_: maximum molar binding ratio
ATA: aurintricarboxylic acid
M-ABB: MOPS-actin binding buffer
DMSO: dimethylsulphoxide
DMF: dimethylformamide
BSA: bovine serum albumin
DTT: dithiothreitol
TCEP: tris(2-carboxyethyl)phosphine
PKA: protein kinase A
FMAL: fluorescein-5-maleimide
TMR: tetramethylrhodamine-5-iodoacetamide
SD: standard deviation
SE: standard errors
DWR: direct waveform recording
MOA: mechanism of action

## REFERENCES CITED

1. Ramaraj, R. (2008) Hypertrophic cardiomyopathy: etiology, diagnosis, and treatment. Cardiol Rev 16, 172–180

2. Refaat, M. M., Fahed, A. C., Hassanieh, S., Hotait, M., Arabi, M., Skouri, H., Seidman, J. G., Seidman, C. E., Bitar, F. F., and Nemer, G. (2016) The Muscle-Bound Heart. Card Electrophysiol Clin 8, 223–231

3. Arad, M., Seidman, J. G., and Seidman, C. E. (2002) Phenotypic diversity in hypertrophic cardiomyopathy. Hum Mol Genet 11, 2499–2506

4. Marsiglia, J. D., and Pereira, A. C. (2014) Hypertrophic cardiomyopathy: how do mutations lead to disease? Arq Bras Cardiol 102, 295–304

5. Maron, B. J., Towbin, J. A., Thiene, G., Antzelevitch, C., Corrado, D., Arnett, D., Moss, A. J., Seidman, C. E., Young, J. B., American Heart, A., Council on Clinical Cardiology, H. F., Transplantation, C., Quality of, C., Outcomes, R., Functional, G., Translational Biology Interdisciplinary Working, G., Council on, E., and Prevention. (2006) Contemporary definitions and classification of the cardiomyopathies: an American Heart Association Scientific Statement from the Council on Clinical Cardiology, Heart Failure and Transplantation Committee; Quality of Care and Outcomes Research and Functional Genomics and Translational Biology Interdisciplinary Working Groups; and Council on Epidemiology and Prevention. Circulation 113, 1807–1816

6. Bozkurt, B., Colvin, M., Cook, J., Cooper, L. T., Deswal, A., Fonarow, G. C., Francis, G. S., Lenihan, D., Lewis, E. F., McNamara, D. M., Pahl, E., Vasan, R. S., Ramasubbu, K., Rasmusson, K., Towbin, J. A., Yancy, C., American Heart Association Committee on Heart, F., Transplantation of the Council on Clinical, C., Council on Cardiovascular Disease in the, Y., Council on, C., Stroke, N., Council on, E., Prevention, Council on Quality of, C., and Outcomes, R. (2016) Current Diagnostic and Treatment Strategies for Specific Dilated Cardiomyopathies: A Scientific Statement From the American Heart Association. Circulation 134, e579–e646

7. Colson, B. A., Thompson, A. R., Espinoza-Fonseca, L. M., and Thomas, D. D. (2016) Site-directed spectroscopy of cardiac myosin-binding protein C reveals effects of phosphorylation on protein structural dynamics. Proc Natl Acad Sci U S A 113, 3233–3238

8. Previs, M. J., Mun, J. Y., Michalek, A. J., Previs, S. B., Gulick, J., Robbins, J., Warshaw, D. M., and Craig, R. (2016) Phosphorylation and calcium antagonistically tune myosin-binding protein C’s structure and function. Proc Natl Acad Sci U S A 113, 3239–3244

9. Jacques, A. M., Copeland, O., Messer, A. E., Gallon, C. E., King, K., McKenna, W. J., Tsang, V. T., and Marston, S. B. (2008) Myosin binding protein C phosphorylation in normal, hypertrophic and failing human heart muscle. J Mol Cell Cardiol 45, 209–216

10. Copeland, O., Sadayappan, S., Messer, A. E., Steinen, G. J., van der Velden, J., and Marston, S. B. (2010) Analysis of cardiac myosin binding protein-C phosphorylation in human heart muscle. Journal of molecular and cellular cardiology 49, 1003–1011

11. Malik, F. I., Hartman, J. J., Elias, K. A., Morgan, B. P., Rodriguez, H., Brejc, K., Anderson, R. L., Sueoka, S. H., Lee, K. H., Finer, J. T., Sakowicz, R., Baliga, R., Cox, D. R., Garard, M., Godinez, G., Kawas, R., Kraynack, E., Lenzi, D., Lu, P. P., Muci, A., Niu, C., Qian, X., Pierce, D. W., Pokrovskii, M., Suehiro, I., Sylvester, S., Tochimoto, T., Valdez, C., Wang, W., Katori, T., Kass, D. A., Shen, Y. T., Vatner, S. F., and Morgans, D. J. (2011) Cardiac myosin activation: a potential therapeutic approach for systolic heart failure. Science 331, 1439–1443

12. Green, E. M., Wakimoto, H., Anderson, R. L., Evanchik, M. J., Gorham, J. M., Harrison, B. C., Henze, M., Kawas, R., Oslob, J. D., Rodriguez, H. M., Song, Y., Wan, W., Leinwand, L. A., Spudich, J. A., McDowell, R. S., Seidman, J. G., and Seidman, C. E. (2016) A smallmolecule inhibitor of sarcomere contractility suppresses hypertrophic cardiomyopathy in mice. Science 351, 617–621

13. Bunch, T. A., Guhathakurta, P., Lepak, V. C., Thompson, A. R., Kanassatega, R. S., Wilson, A., Thomas, D. D., and Colson, B. A. (2021) Cardiac myosin-binding protein C interaction with actin is inhibited by compounds identified in a high-throughput fluorescence lifetime screen. J Biol Chem 297, 100840

14. Schaaf, T. M., Peterson, K. C., Grant, B. D., Bawaskar, P., Yuen, S., Li, J., Muretta, J. M., Gillispie, G. D., and Thomas, D. D. (2017) High-Throughput Spectral and Lifetime-Based FRET Screening in Living Cells to Identify Small-Molecule Effectors of SERCA. SLAS Discov 22, 262–273

15. Bunch, T. A., Lepak, V.C., Bortz, K.M., and Colson, B.A. (2021) A high-throughput fluorescence lifetime-based assay for detecting binding of myosin-binding protein C to Factin. J Gen Physiol

16. Zhang, J. H., Chung, T. D., and Oldenburg, K. R. (1999) A Simple Statistical Parameter for Use in Evaluation and Validation of High Throughput Screening Assays. J Biomol Screen 4, 67–73

17. Guhathakurta, P., Phung, L. A., Prochniewicz, E., Lichtenberger, S., Wilson, A., and Thomas, D. D. (2020) Actin-binding compounds, previously discovered by FRET-based high-throughput screening, differentially affect skeletal and cardiac muscle. J Biol Chem 295, 14100–14110

18. Collibee, S. E., Bergnes, G., Muci, A., Browne, W. F. t., Garard, M., Hinken, A. C., Russell, A. J., Suehiro, I., Hartman, J., Kawas, R., Lu, P. P., Lee, K. H., Marquez, D., Tomlinson, M., Xu, D., Kennedy, A., Hwee, D., Schaletzky, J., Leung, K., Malik, F. I., Morgans, D. J., Jr., and Morgan, B. P. (2018) Discovery of Tirasemtiv, the First Direct Fast Skeletal Muscle Troponin Activator. ACS Med Chem Lett 9, 354–358

19. Kanassatega, R. S., Bunch, T. A., Lepak, V. C., Wang, C., and Colson, B. A. (2022) Human cardiac myosin-binding protein C phosphorylation- and mutation-dependent structural dynamics monitored by time-resolved FRET. J Mol Cell Cardiol 166, 116–126

20. Bunch, T. A., Lepak, V. C., Kanassatega, R. S., and Colson, B. A. (2018) N-terminal extension in cardiac myosin-binding protein C regulates myofilament binding. J Mol Cell Cardiol 125, 140–148

21. Bunch, T. A., Kanassatega, R. S., Lepak, V. C., and Colson, B. A. (2019) Human cardiac myosin-binding protein C restricts actin structural dynamics in a cooperative and phosphorylation-sensitive manner. J Biol Chem

22. Gruber, S. J., Cornea, R. L., Li, J., Peterson, K. C., Schaaf, T. M., Gillispie, G. D., Dahl, R., Zsebo, K. M., Robia, S. L., and Thomas, D. D. (2014) Discovery of enzyme modulators via high-throughput time-resolved FRET in living cells. J Biomol Screen 19, 215–222

23. Schaaf, T. M., Peterson, K. C., Grant, B. D., Thomas, D. D., and Gillispie, G. D. (2017) Spectral Unmixing Plate Reader: High-Throughput, High-Precision FRET Assays in Living Cells. SLAS Discov 22, 250–261

24. Talbot, C. B., Lagarto, J., Warren, S., Neil, M. A., French, P. M., and Dunsby, C. (2015) Correction Approach for Delta Function Convolution Model Fitting of Fluorescence Decay Data in the Case of a Monoexponential Reference Fluorophore. J Fluoresc 25, 1169–1182

25. Guhathakurta, P., Rebbeck, R. T., Denha, S. A., Keller, A. R., Carter, A. L., Atang, A. E., Svensson, B., Thomas, D. D., Hays, T. S., and Avery, A. W. (2023) Early-phase drug discovery of beta-III-spectrin actin-binding modulators for treatment of spinocerebellar ataxia type 5. J Biol Chem 299, 102956

26. Rebbeck, R. T., Essawy, M. M., Nitu, F. R., Grant, B. D., Gillispie, G. D., Thomas, D. D., Bers, D. M., and Cornea, R. L. (2017) High-Throughput Screens to Discover Small-Molecule Modulators of Ryanodine Receptor Calcium Release Channels. SLAS Discov 22, 176–186

27. Goutelle, S., Maurin, M., Rougier, F., Barbaut, X., Bourguignon, L., Ducher, M., and Maire, P. (2008) The Hill equation: a review of its capabilities in pharmacological modelling. Fundam Clin Pharmacol 22, 633–648

28. Chen, X., Tume, R. K., Xu, X., and Zhou, G. (2017) Solubilization of myofibrillar proteins in water or low ionic strength media: classical techniques, basic principles, and novel functionalities. Crit Rev Food Sci Nutr 57, 3260–3280

29. Ma, Y. Z., and Taylor, E. W. (1994) Kinetic mechanism of myofibril ATPase. Biophys J 66, 1542–1553.

30. Smith, S. H., and Fuchs, F. (1999) Effect of ionic strength on length-dependent Ca(2+) activation in skinned cardiac muscle. J Mol Cell Cardiol 31, 2115–2125

31. Bers, D. M., Patton, C. W., and Nuccitelli, R. (2010) A practical guide to the preparation of Ca(2+) buffers. Methods Cell Biol 99, 1–26

