## Supporting information for "Drug discovery for heart failure targeting myosin-binding protein C"

*Equal contribution

‡Co-senior authors

†Corresponding authors

Running title: *High-throughput assay detecting actin-cMyBP-C modulators*

**Keywords:** actin, cardiac muscle, cardiac myosin-binding protein C (cMyBP-C), contractile proteins, phosphorylation, protein kinase A (PKA), fluorescence lifetime (FLT), high throughput screen (HTS), fluorescence resonance energy transfer (FRET), site-directed spectroscopy

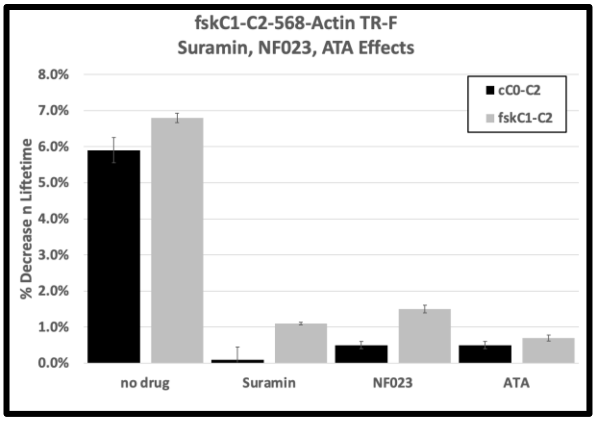


Figure S1. Suramin, NF023 and ATA inhibit fskC1-C2 binding to actin. Suramin (20 μM), NF023 (50 μM) and ATA (20 μM) decreased the binding of fskC1-C2 (2 μM, grey) to 568-Actin (1 μM). The effects on cC0-C2 (black) as described in (13) are shown for comparison. Values are the % decrease in FLT of AF568-actin due to cC0-C2 or fskC1-C2, and are the average ± SE. n=3-5.

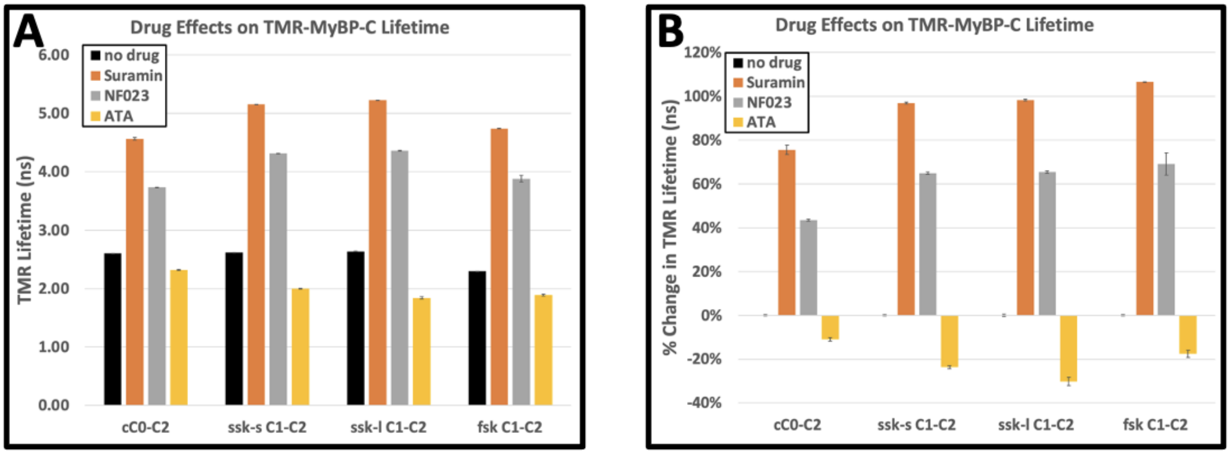


**Figure S2.** Suramin, NF023 and ATA change the FLT of TMR on cardiac C0-C2 and skeletal C1-C2. cC0-C2, ssk-sC1-C2, ssk-lC1-C2, and fskC1-C2 were labeled with TMR whose FLT in the absence of compound was 2.30-2.64 nm (black). Suramin (20 μM, orange) and NF023 (50 μM, grey) increased the FLT of TMR, while ATA (20 μM, gold) decreased its FLT on all isoforms. **A.** FLT in ns is shown. **B.** The percent change in FLT compared to no compound is displayed. All values are the average ± SE. n=5.

**Table S1.** FMAL-actin lifetime changes with TMR-C0-C2 binding

| Screen | | Average lifetime (ns) | SD | CV (%) | n | Change FRET (%) | Z’ |
|---|---|---|---|---|---|---|---|
| 1 | NoTMR-C0-C2 | 4.11 | 0.02 | 0.59 | 624 | | |
| | WithTMR-C0-C2 | 3.54 | 0.05 | 1.39 | 624 | 13.8 | 0.61 |
| 2 | No TMR-C0-C2 | 3.90 | 0.02 | 0.57 | 624 | | |
| | With TMR-C0-C2 | 3.45 | 0.03 | 0.95 | 624 | 11.4 | 0.63 |

Experiments were carried out with two separate protein preparations for FMAL-F-actin (0.25 µM, 10% labeled) without and with 2 separate TMR-cC0-C2 preparations (0.5 µM, 70-90% labeled). The average FLT of FMAL, attached to F-actin (in ns), standard deviation (SD), number of wells (n), change in FLT (FRET) due to TMR-cC0-C2, and Z´ factor are shown [N=2, n=1248].

**Table S2 Reproducible hits tested in CRC (continued in next page)**

| Compounds | Z' score<br>Screen 1 FRET | Z' score<br>Screen 2FRET | Z' score<br>Screen 1 FLT | Z' score<br>Screen 2 FLT |
|---|---|---|---|---|
| 1. Ampicillin sodium | -8.28 | -8.02 | 1.77 | 1.26 |
| 2.. Anidulafungin | -0.32 | -0.99 | 17.34 | 16.03 |
| 3. Bedaquiline fumarate | 15.82 | 7.16 | -3.21 | -2.25 |
| 4. Benzonatate | -4.99 | -1.23 | -5.30 | -3.83 |
| 5. Cefepime Dihydrochloride Monohydrate | -6.07 | -5.45 | -2.80 | 0.55 |
| 6. Cefsulodin sodium | -19.27 | -16.06 | 5.15 | 1.58 |
| 7. Chloroquine Phosphate | 0.95 | 1.56 | 11.03 | 7.59 |
| 8. Clindamycin palmitate HCl | 19.91 | 14.78 | -2.61 | -2.63 |
| 9. Clinofibrate | -4.35 | -3.23 | 0.88 | 1.15 |
| 10. Diammonium Glycyrrhizinate | -3.96 | -3.85 | 0.96 | 0.95 |
| 11. Doxazosin | | | | |
| 12. Elbasvir | -4.08 | -3.24 | 0.27 | 0.39 |
| 13. Eltrombopag Olamine | | | | |
| 14. Ertapenem sodium | -9.20 | -9.56 | 4.64 | 3.73 |
| 15..Erlotinib | -0.82 | -0.64 | 4.90 | 3.56 |
| 16.Erlotinib HCl | | | | |
| 17. Erythromycin estolate | -4.08 | | -0.32 | |
| 18.Enoxaparin sodium | -6.76 | -4.12 | 0.45 | 0.51 |
| 19. Febuxostat | -1.58 | -2.03 | 6.02 | 5.48 |
| 20. Fenticonazole Nitrate | 5.27 | 3.08 | -0.86 | 0.11 |
| 21. Fidaxomicin | -4.67 | -3.16 | -0.81 | 0.21 |
| 22.Flopropione | | | | |
| 23. Fostamatinib (R788) | -5.64 | -3.15 | -0.70 | -3.21 |
| 24..Hederagenin | -3.93 | -3.15 | -0.71 | -0.82 |
| 25. Heparin sodium | -11.06 | -10.68 | -0.47 | -1.29 |
| 26. Hyodeoxycholic acid (HDCA) | | | | |
| 27. Laquinimod | -4.34 | -3.71 | 0.38 | 0.72 |
| 28. Latamoxef sodium | -5.87 | -3.38 | -2.60 | -0.95 |
| 29.Lumiracoxib | -6.57 | -4.57 | -1.92 | -0.71 |
| 30. Luteolin | | | | |
| 31. Masitinib (AB1010) | | | | |
| 32. Nilotinib hydrochloride | 1.83 | 0.80 | 5.98 | 4.04 |
| 33. Olsalazine Sodium | -11.93 | -6.49 | -1.52 | -4.39 |
| 34. Oxytetracycline (Terramycin) | -8.44 | -6.40 | -0.42 | -1.74 |
| 35. Paritaprevir (ABT-450) | -1.56 | -1.59 | 10.33 | 7.94 |
| 36.Pneumocandin B0 | 1.47 | 1.75 | 8.34 | 5.20 |
| 37. Pranlukast | -0.62 | -2.81 | 4.25 | 4.04 |
| 38.. Proanthocyanidins | | | | |
| 39.. Proflavine Hemisulfate | | | | |
| 40. Protoporphyrin IX | -6.12 | -4.33 | 0.88 | 0.04 |
| 41.. Pyrantel Pamoate | | | | |
| 42. Raloxifene | | | | |
| 43. Regorafenib HCl | | | | |
| 44. Regorafenib (BAY 73-4506) | -0.30 | | 5.17 | |
| 45. Relugolix | | | | |
| 46. Sacubitril/valsartan (LCZ696) | -6.40 | -3.33 | 1.16 | 0.98 |
| 47. Saikosaponin D | 3.48 | 3.38 | 4.00 | 3.51 |
| 48.. Sanguinarine chloride | | | | |
| 49.Scutellarin | -6.84 | -3.74 | -3.89 | -4.16 |
| 50. Tamibarotene | -5.18 | -4.18 | 3.71 | 3.19 |
| 51.Telotristat Etiprate (LX 1606 Hippurate) | | | | |
| 52. Thonzylamine | | | | |
| 53. Tocofersolan | 6.00 | 4.88 | 3.00 | 4.14 |
| 54. Triclocarban | 3.16 | 1.43 | 15.72 | 11.50 |
| Table S2 continued | | | | |
| 55. Troxerutin | 1.31 | 0.81 | 4.01 | 3.60 |
| 56. Valsartan | -6.40 | -3.30 | 1.16 | 0.98 |

| | | | | |
|---|---|---|---|---|
| 57. Verteporfin* | -3.63 | | 3.14 | |
| 58.Zaltoprofen | -6.37 | -3.83 | -0.36 | 1.10 |
| 59. α-Hederin | 1.59 | 3.33 | 4.94 | 4.77 |
| 60.8-Hydroxyquinoline | 0.43 | 6.94 | -6.52 | -7.07 |

Z’ scores were not shown for the compounds, which were specifically identified in approach 2. See experimental procedure
